# Severe childhood and adulthood stress associates with neocortical layer-specific reductions of mature spines in psychiatric disorders

**DOI:** 10.1101/2020.08.31.276386

**Authors:** Dominic Kaul, Caine C Smith, Julia Stevens, Anna S Fröhlich, Elisabeth B Binder, Naguib Mechawar, Sibylle G Schwab, Natalie Matosin

## Abstract

Severe stress exposure causes the loss of dendritic spines on cortical pyramidal neurons and induces psychiatric-like symptoms in rodent models. These effects are strongest following early-life stress and are most persistent on apical dendrites. However, the long-term impacts and temporal effects of stress exposure on the human brain remain poorly understood. Using a novel postmortem cohort of psychiatric cases with severe stress experienced in childhood, adulthood, or no severe stress, and matched controls, we aimed to determine the impact of stress timing on pyramidal neuron structure in the human orbitofrontal cortex (OFC). We performed Golgi Cox staining and manually measured the morphology and density of over 22,000 dendritic spines on layer-specific pyramidal neuron apical dendrites. We also quantified glucocorticoid receptor mRNA and protein as a marker of stress dysregulation. Both childhood and adulthood stress were associated with large reductions in mature mushroom spine density (up to 56% loss) in both the superficial (II/III) and deeper layers (V) of the OFC. However, childhood stress caused more substantial reductions to both total and mature mushroom spines. No difference in glucocorticoid receptor mRNA and protein were seen between groups, although both negatively correlated with total spine density within the whole cohort. These findings indicate that severe stress, especially when experienced during childhood, persistently affects the fine morphological properties of neurons in the human OFC. This may impact on cell connectivity in this brain area, and at least partly explain the social and emotional symptoms that originate in the OFC in psychiatric disorders.

## 1. Introduction

Severe stress is among the most well-supported environmental factors involved in the development of psychiatric disorders, particularly early-life stress [1]. These disorders include depression, bipolar disorder, and schizophrenia, three of the highest burden mental illnesses globally [2]. Their shared neuropathology and symptoms often make categorical diagnosis difficult and determining the contributions of stress towards their development has been complicated by several factors. Conditional experiences, such as duration and timing of stress exposure may differentially determine how an individual is affected [3, 4], with repeated and chronic exposure adding to the diagnostic complexity of these conditions [5]. Biological subtypes of psychiatric disorders based on environmental exposures likely exist and elucidating these subtypes may be a pre-requisite for developing more personalised treatment interventions [6].

The orbitofrontal cortex (OFC) is closely involved in stress regulation. It is important for emotion-related learning and evaluating reinforcers of behaviour [7], integral to the perception and response to stress. Reduced OFC connectivity with both cortical and subcortical regions has been associated with altered fear responses and social anxiety [8], Moreover, lesions in the non-human primate OFC cause impaired emotional regulation (e.g. anxiety [9]) and social behaviour [10], symptoms seen in psychiatric disorders. Early-life psychosocial stress has been associated with reductions of OFC volume which correlated with social difficulties in later life [11]. Reduced OFC volume is also repeatedly identified in psychiatric disorders and is associated with symptom severity [12, 13]. However, imaging studies do not provide the resolution to determine the underlying molecular and cellular changes. In adult rats, chronic stress causes persistent transcriptional modifications [14] and physical cellular alterations in the OFC including the retraction of glutamatergic neuron processes [15, 16], reductions in the density of GABAergic neurons [17], as well as alterations to glial activity and connectivity [18]. Whether similar changes also occur in the human OFC after stress is poorly understood. This remains pertinent as the rodent OFC is agranular [19], and is not homologous to neither the non-human primate OFC nor the human OFC [19, 20]. Consequently, whilst we consider OFC of rodents, it must be recognised that the effects noted in this region may not necessarily translate into humans, further incentivising study of human cohorts.

Pyramidal neurons are integral components of neural circuitry that exhibit high numbers of dendritic spines [21]. Dendritic spines are 1-10μm protrusions from neuronal dendrites that are sites of several essential postsynaptic proteins, neurotransmitter, neurotrophic, and hormone receptors [22]. The unique structure, localisation and motility of dendritic spines play a role in the compartmentalisation of neuronal signalling [23] making these structures key indicators of neural connectivity. Dendritic spine morphology is integral to synaptic functionality [24], with spine head size positively associated with receptor density, spine stability and spine motility [25].Both spine distribution and morphology are essential to neural network function.

Rodent models consistently demonstrate that stress reduces pyramidal neuron dendrite complexity, spine density, and spine head size in the OFC [16, 26–28]. When stress is experienced during early-life, reduced spine density can persist into adulthood on apical dendrites [27]. In rodent stress models, loss of cortical dendritic spines have also been associated with depression-related phenotypes [29]. In humans, reductions in post-synaptic proteins in the OFC have been observed in mood disorders and schizophrenia [30, 31], suggesting dendritic spines may be involved in impairments of OFC functions associated with these disorders. In one human postmortem study, reduced total and mature dendritic spine density in the OFC were associated with increased expression of stress hormone regulators associated with early-life adversity indicating that spines in the human OFC are sensitive to stress [32]. However, the temporal effects of stress on dendritic spines in the human brain and their contribution towards psychopathology are unknown.

Stress, particularly early-life stress, causes persistent dysregulation of the glucocorticoid receptor (GR) [33]. Long-term reprogramming of GR can impact neural connectivity and contribute towards the development of psychiatric disorders [34, 35]. A complete knockdown of GR in the prefrontal cortex (PFC) of Sprague-Dawley rats induces stress hypersensitivity and depression-like behaviour [36]. Cortical GR knockdown also attenuates losses of apical dendritic spines in adult female mice [37]. In pyramidal neurons, extranuclear GR is localised to the postsynaptic density (PSD) of dendritic spines [38, 39] and increased activation of GR increases spineogenesis and maturation [40] while inhibition significantly reduces spineogenesis, even when compensated with artificially increased corticosteroids [41].

Persistent alterations to GR expression may influence the distribution of dendritic spines and may be important to remediating stress-induced alterations to dendritic spines.

The overlap between the impacts of stress on dendritic spines and their neuropathology in severe psychopathology indicate a shared cytoarchitectural pathway. We aimed to better understand this relationship. We comprehensively examined the influence of severe childhood and adulthood stress on dendritic spines and GR alterations in the postmortem human OFC, across the spectrum of severe psychotic and mood disorders. Our findings support that severe stress exposure causes persistent cytoarchitectural remodelling in the human OFC, which might be intrinsic to the development and symptomatology of psychiatric disorders.

## 2. Methods

### 2.1. Postmortem cohort selection

Ethical approval for this study was obtained from the Human Ethics Committee at the University of Wollongong (HE2018/351). Postmortem brain tissue was collected at the NSW Brain Tissue Resource Centre (Sydney, Australia). Informed consent was given by all donors or their next of kin for brain autopsy. History of stressful events (childhood/adulthood) and psychiatric diagnoses (schizophrenia/schizoaffective disorder/depression/bipolar disorder) were extracted from extensive medical records of 103 subjects. Severe stressors were considered to be any event, series of events, or set of circumstances that was physically or emotionally harmful or threatening, and had lasting adverse effects on the individual’s functioning and physical, social, or emotional well-being (e.g. physical/emotional abuse or neglect). Of the original 103 cases, 32 subjects with the most applicable stress histories were selected constituting four groups: severe childhood stress with psychopathology, severe adulthood stress with psychopathology, psychopathology with no severe stress, and healthy controls with no severe stress exposure or psychopathology (Supplementary Table 1; *n*=8 per group). Groups were matched according to psychiatric diagnoses, postmortem interval (PMI), age at death, and RNA integrity number (RIN) (Table 1). All experiments were performed blind with coded subjects.

**Table 1.** Summary of demographic and clinical variables for the cohort. Values are presented as the mean ± SEM.

|  | Childhood stress and psychiatric disorder | Adult stress and psychiatric disorder | No severe stress and psychiatric disorder | Control (no stress, no psychiatric disorder) | ANOVA<sup>1</sup> (F<sub>3,28</sub>) |
| --- | --- | --- | --- | --- | --- |
| **Sample size** | 8 | 8 | 8 | 8 |  |
| **Age (Years)** | 59.5 ± 7.2 | 52.4 ± 4.5 | 50.8 ± 3.9 | 55.1 ± 5.5 | 0.49618<br>P=0.687886 |
| **Sex (M/F)** | 6/2 | 6/2 | 5/3 | 5/3 | X<sup>2</sup>=0.5818<br>P=0.9906 |
| **PMI (Hours)** | 35.3 ± 6.2 | 45.8 ± 6.4 | 31.6 ± 3.4 | 32.2 ± 3.6 | 1.65922<br>P= 0.19837 |
| **Brain pH** | 6.47 ± 0.09 | 6.50 ± 0.10 | 6.52 ± 0.05 | 6.77 ± 0.04 | **3.40475**<br>**P=0.031249** |
| **RIN** | 6.44 ± 0.50 | 6.86 ± 0.66 | 7.35 ± 0.27 | 7.98 ± 0.20 | 2.1695<br>P=0.11391 |
| **Number of cases** | 5SZ,2MDD,1SZA | 4SZ,1MDD,2BD,1SZA | 5SZ,2MDD,1BD | - | $\chi^2= 0.3429$<sup>2</sup><br>P=0. 8425 |

<sup>1</sup>Unless otherwise stated
<sup>2</sup>Performed as number of mood disorder cases (MDD, BD, SZA) and psychotic disorder cases (SCZ)
Abbreviations: M=male, F=female, PMI=postmortem interval, RIN=RNA integrity number, SZ=schizophrenia, MDD=major depressive disorder, BD=bipolar disorder, SZA=schizoaffective disorder

### 2.2. Golgi-Cox staining

Fresh frozen tissue blocks (~0.5cm^3^) from the OFC were stained using the FD Rapid Golgistain™ Kit following the manufacturer’s instructions (FD Neurotechnologies, Columbia, MD, USA). Coronal sections (150μm) were cut from frozen tissue blocks. Sections were transferred onto gelatin-coated microscope slides and completely dried in a vacuum desiccator at room temperature (RT) for 24-48 hours. Sections were rehydrated on slides in water and then gradually dehydrated of developing solution by immersions in increasing concentrations of ethanol. Washed and dehydrated sections were cleared in xylene and coverslipped. Sections were stored in the dark at RT until imaging.

### 2.3. Dendritic spine sampling

All microscopy was completed using brightfield imaging (DMi8; Leica Microsystems, Wetzlar, Germany). At 10x magnification, regular stratified cortical layers (I-VI) were defined according to previously outlined morphological features (Figure 1a) [42]. Pyramidal neurons are classically found in layers II, III and V, which were thus selected for analysis [42]. Layers II and III were difficult to distinguish under the Golgi-Cox stain and thus combined for analysis. For each section, an adapted optical fractionator method was used to randomly select coordinates within each layer. This was repeated until 3-6 pyramidal neurons were imaged. Neurons with poor staining quality were rejected and new neurons selected. Z-stacks of selected neurons centred on apical dendrites were collected at 63x magnification in oil using a 0.6μm step-size at a resolution of 2048 x 2048 pixels (Figure 1b/c). Settings for exposure and gain were optimised and kept constant between samples.

**Figure 1.**
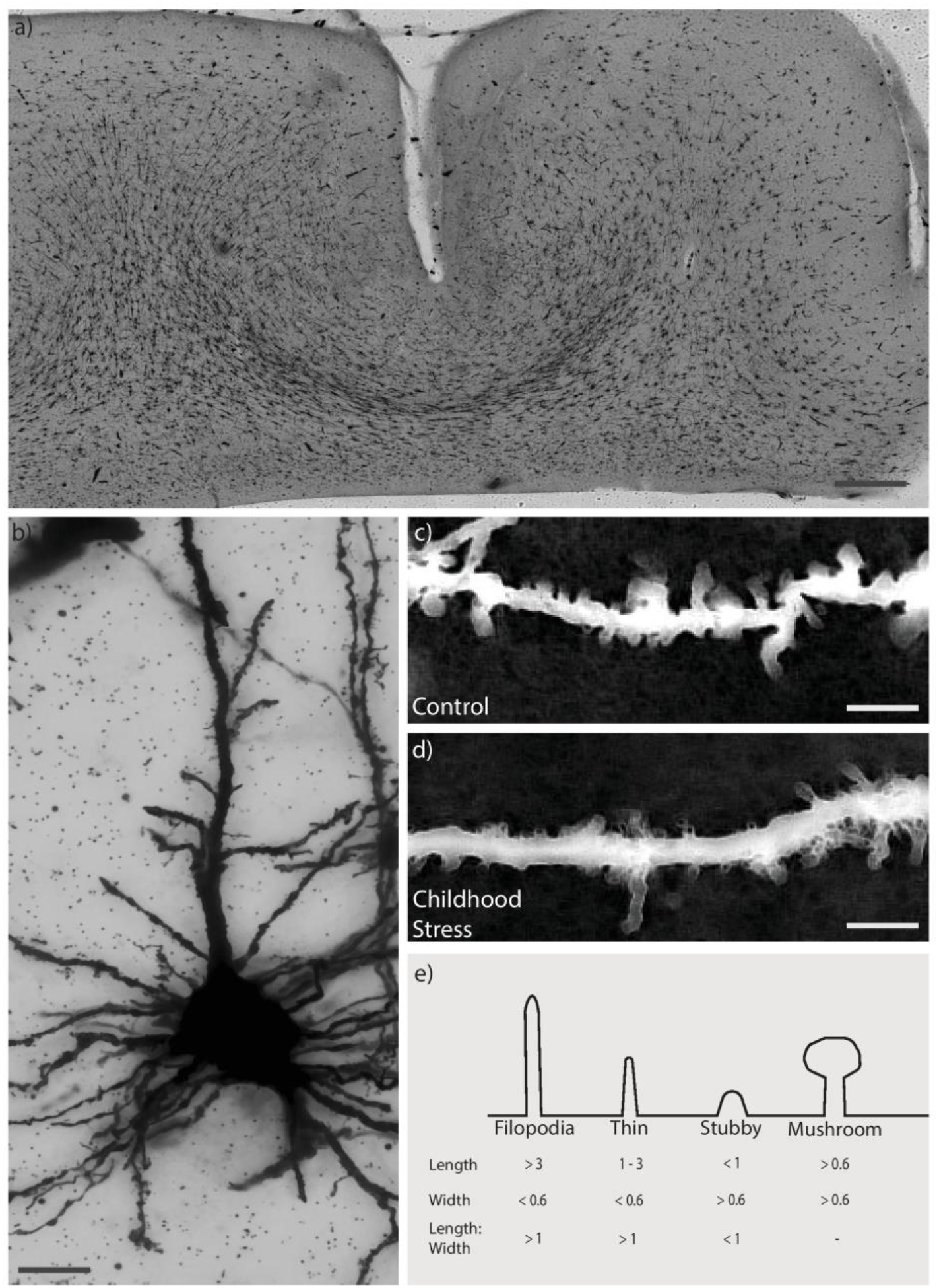
Representative OFC section, neuron and spines used for analysis. (**a)** Tilescan of Golgi-Cox stained section (150μm thick). Scale bar indicates 1mm. (**b)** Representative merge of pyramidal neuron from z stack using minimum thresholding. Scale bar indicates 10μm. (**c/d)** Representative segment of apical dendritic process, inverted and merged using maximum thresholding in childhood stress and control group Scale bar indicates 5μm. **(e)** Dendritic spines were classified into four distinct groups based on measurements of spine length (base to tip) as well as width (at the widest point)

Spine sampling was conducted as outlined previously [43, 44]. Selected dendritic segments were <3μm in diameter, within 100μm of the coronal surface, relatively parallel to the surface of the section, separate from other dendritic segments, and >20μm in length. Along the apical dendrite of each accepted neuron, 3-6 randomly selected segments that met these criteria were analysed using ImageJ with the Fiji plugin [45]. Dendrites were traced to determine segment length. All dendritic spines located along the segment within the same x-y plane as the segment were counted, measured manually and morphologically categorised as outlined previously [46] with slight modifications based on trial sampling (Figure 1d).

### 2.4. Quantitative PCR

Total RNA was extracted using the RNeasy Plus Universal Mini Kit following the manufacturer’s instructions (QIAGEN, Hilden, Germany). cDNA was synthesised from total RNA using the High Capacity RNA-to-cDNA Kit (Applied Biosystems, Waltham, MA, USA). TaqMan gene expression assays (Applied Biosystems) were used to quantify *NR3C1* (Hs00353740_m1) and housekeeper genes ACTB (Hs99999903_m1) and GAPDH (Hs99999905_m1). Per replicate, 50 ng of total cDNA was combined with TaqMan Master Mix (Applied Biosystems) and TaqMan assay and quantified using the QuantStudio5 PCR system (Thermo Scientific, Waltham, MA, USA). Reactions were performed in triplicate. Gene expression was normalised against the geometric mean of ACTB/GAPDH. Outputs were analysed by the 2^-ΔΔCt^ method[47], relative to the control group.

### 2.5. Western blot

Protein was extracted and quantified using Bradford Protein Assay (Bio-Rad, Hercules, CA, USA). 20μg total protein was separated using SDS-PAGE and transferred to PVDF membranes. Blots were blocked in a Tris-buffered saline with 0.1% Tween 20 (TBST) and 5% skim milk for one hour at RT, then incubated overnight in primary antibodies for GR (1:1,000; sc-393232, SantaCruz Biotechnology, Santa Cruz, CA, USA) and β-actin (1:10,000; a1978-200UL, Sigma-Aldrich, St. Louis, MO, USA), diluted in TBST with 1% skim milk. Blots were washed and probed with secondary antibody for one hour at RT (anti-mouse IgG, horseradish peroxidase linked; 1:5,000; Merck, Burlington, MA, USA). Bands were visualized using high sensitivity Pierce ECL Plus (Thermo Scientific) and quantified using the Amersham Imager 600 (GE Healthcare, Chicago, IL, USA). The optical density of GR was normalised to β-actin and a cross-membrane control.

### 2.6. Statistical analysis

Statistics were performed in R. Significance was set to P<0.05 and data presented as mean±SEM. One outlier (>2 mean±SD) was identified for GR mRNA levels (adulthood stress) and was removed. Normality was assessed using Shapiro-Wilks tests. Data not normally distributed were normalised using loge transformation. Analysis of variance (ANOVA) followed by Tukey HSD test post-hoc were performed to compare mean spine and GR measures between groups. One-way analysis of covariance (ANCOVA) accounting for potentially confounding factors determined with Pearson/Spearman correlations or t-tests, for age at death, PMI, RIN, pH, diagnosis, sex (male/female) and psychotic features (yes/no) were subsequently performed with post-hoc pairwise comparisons between groups. Logistic regressions were used to correlate spine densities, segment distance from soma, and GR mRNA/protein.

## 3. Results

### 3.1. Dendritic spines are altered in a morphology-and layer-specific manner and associate with stress history

To determine whether stress exposure impacts on dendritic spine density and morphology, we evaluated apical dendrites of pyramidal neurons in layer II/III and layer V subfields. On average, four segments on four neurons were counted for both subfields in the 32 individuals (1,024 segments). A total of 22 688 dendritic spines (12,144 in Layer II/III, 10,544 from layer V) were counted and classified based on spine morphology. Of the total spine count, filopodia constituted 6.8% (1,556), thin spines 40.1% (9,087), stubby spines 39.6% (8,981), and mushroom spines 13.5% (3,064). Number of spines were normalised to segment length to give a value of spine density/μm. Measurements of spine density for each case were taken as the average of all measured segments.

We initially assessed potential differences in total spine density between stress groups (Figure 2a). There was a significant difference in layer II/III total spine density between groups (F_3,28_=3.06, P=0.045), driven by lower mean spine density in psychiatric disorder cases with childhood stress compared to cases with no stress (−21%; P=0.028). In layer V, there was no significant difference in total dendritic spine density (F_3,28_=0.76, P=0.53; Figure 2a), and after accounting for age (F_3,27_=0.72, P=0.55).

**Figure 2.**
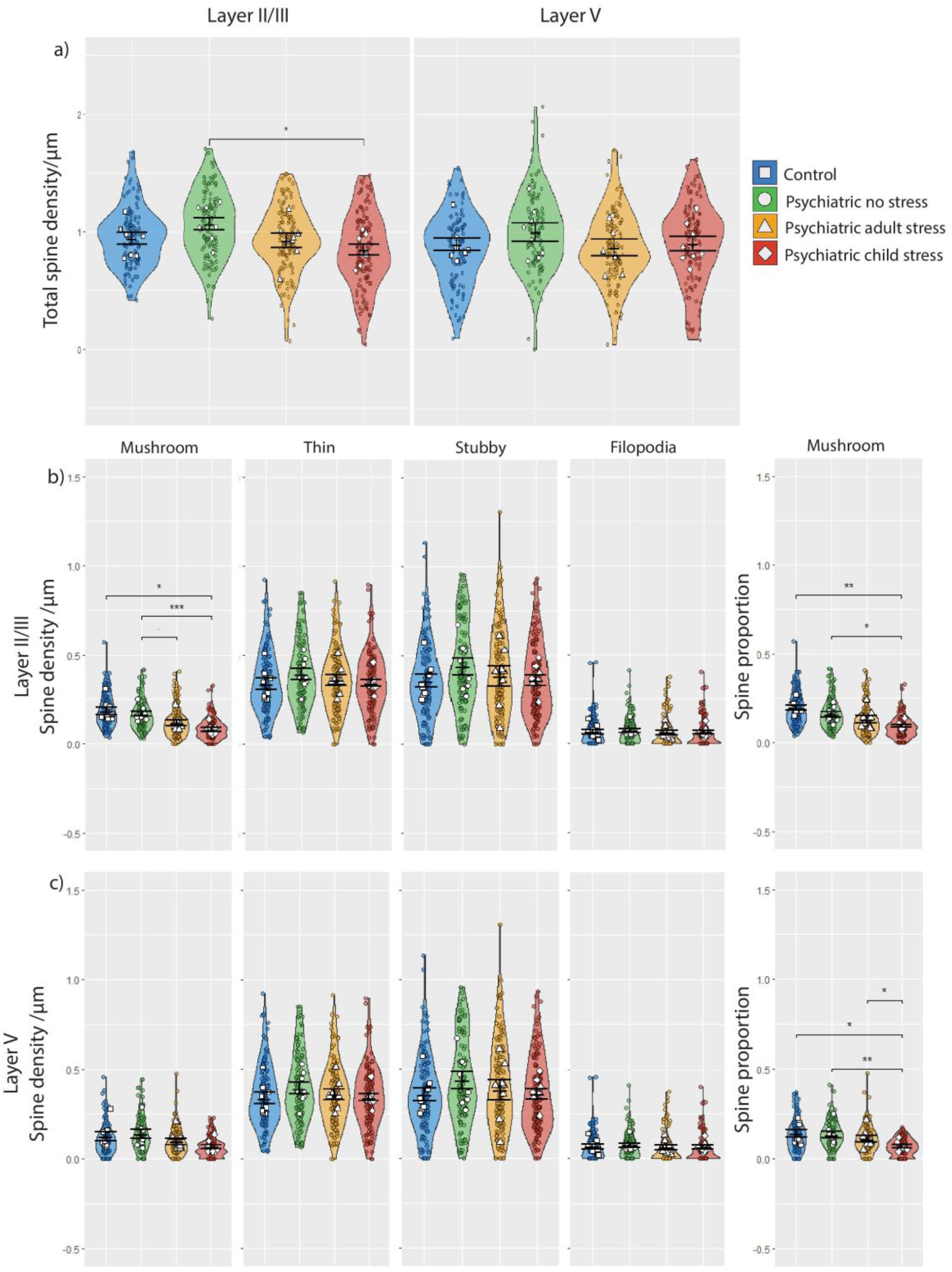
(**a**) Total spine density/μm in layers II/III and layer V across the four stress groups. (**b**) Spine density of each morphologically delineated dendritic spine type and proportion of total spine count in superficial layer II/III. (**c**) Spine density of each morphologically delineated dendritic spine type and proportion of total spine count in deep layer V. Data are presented as means ± SEM. Violin plots and coloured points indicate distribution of individual segments used to calculate case mean. Significance in pairwise comparisons is indicated by * and refers to the results of ANCOVA (*P<0.05, **P<0.01, ***P<0.001).

In layer II/III, severe stress had a significant effect on mushroom spine density (F_3,28_=9.47, P=0.0002). This remained significant after accounting for pH and RIN (F_3,26_=6.412, P=0.002), with lower mushroom spine density in cases with childhood stress compared to both controls (−56%; P=0.016) and cases without stress (−52%; P=0.00099; Figure 2b). This was not seen in cases with adulthood stress, although a trend of reduced spine density was identified, compared to cases without stress (−30%; P=0.059). Mushroom spine density was also altered in layer V (F_3,28_=3.90, P=0.019) due to fewer mushroom spines in cases with childhood stress compared to cases without stress (−53%; P=0.019), although this effect was borderline after correcting for RIN (F_3,27_=2.78, P=0.058; Figure 2c). Thin, stubby and filopodia spines were not altered in layer II/III or V.

To determine proportional changes to spine types, segment density for each spine type was taken over total segment spine density (Figure 2b/c). In layer II/III, stress had a significant effect on the proportion of mushroom spines (F_3,28_=9.01, P=0.0002), with a reduced proportion of mushroom spines in the childhood stress group compared to both controls (−52%; P=0.0001) and cases with no stress history (−40%; P=0.020), as well as the adulthood stress group compared to controls (−33%; P=0.017). These effects persisted after correcting for RIN and pH (F_3,26_=5.218, P=0.006) between the childhood stress and both the control group (P=0.001) and no stress group (P=0.019; Figure 2b). Similarly, mushroom spine proportion in layer V was reduced (F_3,28_=5.39, P=0.005; ANCOVA: F_3,27_=3.51, P=0.029; Figure 2c), in the childhood stress group compared to controls (−51%; ANOVA, P=0.007; ANCOVA, P=0.020), cases with no stress history (−40%; ANOVA, P=0.011; ANCOVA, P=0.004) and cases with adulthood stress compared to controls (ANCOVA, −33%; P=0.039). The proportion of filopodia, stubby, and thin spines in both layer II/III and layer V were not significantly associated with stress exposure.

We next correlated individual segment spine density with the distance from the soma, to determine spatial effects (Figure 3). In the total cohort (*n*=32), total spine density and distance from the soma were negatively correlated in layers II/III and V (Figure 3). Mushroom spine density was also negatively correlated with distance from the soma in layer II/III but not layer V. Thin spines were additionally negatively correlated with segment distance in layer II/III (Supplementary Figure 1). While no overall correlation was observed in thin spines in layer V, there was an interactive effect between the control group and psychiatric cases with no stress history (P=0.0004), and those with adulthood stress history (P=0.0009), indicating that controls displayed a stronger negative correlation. No correlation nor interactive effects were identified for stubby or filopodia spines (Supplementary Figure 1).

**Figure 3.**
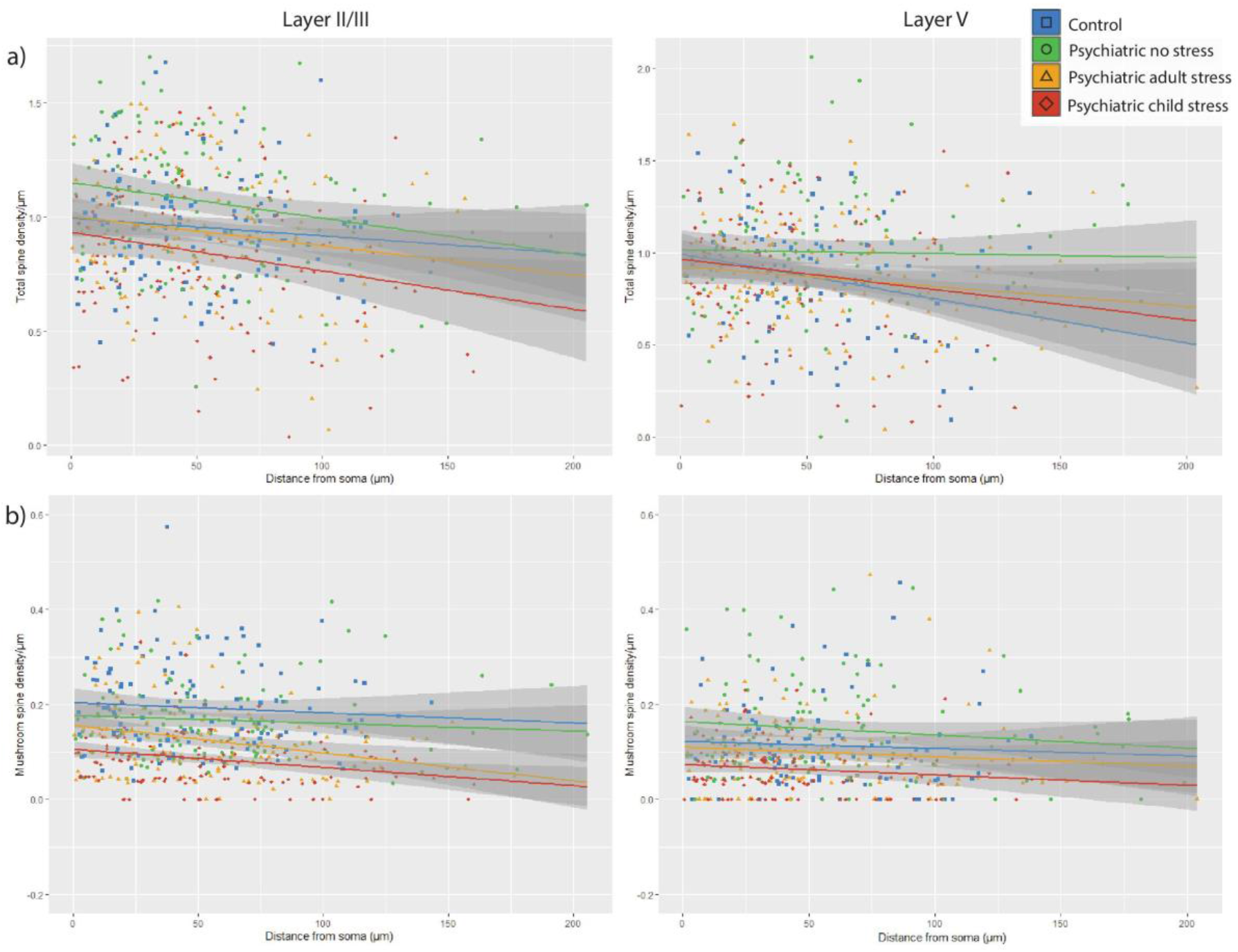
Linear correlations (± confidence interval) (**a**) between segment distance from the soma and segment total spine density. In the whole cohort, significant correlation identified in layers II/III (t=3.836, P=0.0001, R^2^=0.0269) and V (t=2.333, P=0.0201, R^2^=0.0100). (**b**) Between segment distance from the soma and mushroom spine density, measured along the dendritic segment. No interactive effect between groups was identified.

### 3.2. Glucocorticoid receptor expression does not associate with stress history nor spine density in the orbitofrontal cortex

For qRT-PCR, control samples (no template and no reverse transcriptase enzyme) produced no signal for GR and ACTB assays. For western blotting, single bands for GR and β-actin were detected at the expected molecular weights (94 kDa and 42 kDa respectively; Supplementary Figure 2). There was no difference in relative GR mRNA or protein levels between the stress groups (Figure 4a/b). We subsequently examined the relationship between total spine density and GR expression. As GR quantification was performed in bulk tissue, spine density for these correlations were taken as the average of layer II/III and V density per individual (Supplementary Table 2). There was borderline significant correlation between total spine density and GR mRNA expression, and significant correlation with protein when assessing all subjects together. Group-level analyses revealed no correlation between GR mRNA or protein with total spine density, although mushroom spine density and GR mRNA were correlated when assessing all subjects together; this did not hold in individual stress groups, or for protein levels (Figure 4c/4d). We also performed correlations for thin, stubby, and filopodia spine densities and performed exploratory correlation with layer-specific densities. A significant correlation between GR protein and thin spine density was observed within the whole cohort and the adulthood stress group, and stubby spines within the childhood stress group (Supplementary Table 2).

**Figure 4.**
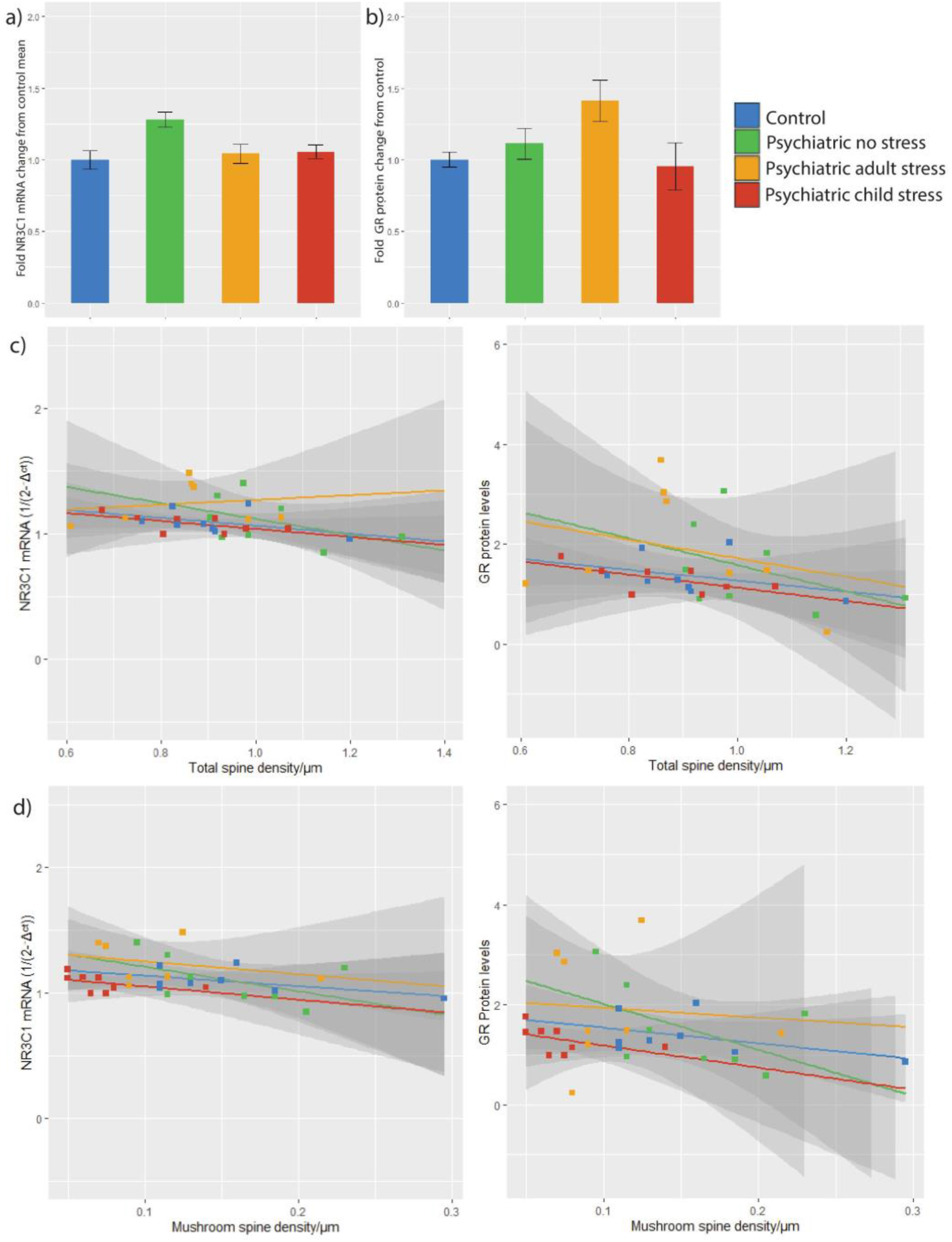
Analysis of glucocorticoid receptor levels in the OFC and the correlation with spine density (**a)** NR3C1 mRNA Ct relative to the geometric mean of ACTB and GAPDH Ct, all reactions performed in triplicate. Data are means ± SEM. F_3,27_=0.921, P=0.443 (**b)** Glucocorticoid receptor protein levels relative to β-actin, normalised across blots, performed in duplicate. F_3,28_=0.169, P=0.916 (**c**) Correlation of GR mRNA levels and total spine densities (average of layer II/III and V density). (**d**) Correlation of GR mRNA levels and mushroom spine densities (average of layer II/III and V density). Data are linear correlations (± confidence interval)

## 4. Discussion

Dendritic spines are repeatedly implicated in psychiatric disorders [48, 49], yet the role of stress in this process remains poorly understood. We provide the first evaluation of dendritic spines in the postmortem OFC derived from individuals with psychiatric disorders. The well-defined temporal stress histories of individuals in this cohort enabled novel comparison between the effects of stress exposure at both childhood and adulthood time points. We identified that psychiatric patients with a history of severe stress, particularly during childhood, display substantial losses in dendritic spines. These losses were most evident in mature mushroom spine populations in the superficial layers of the OFC.

We observed that the number of dendritic spines was reduced in cases with a history of severe stress, and most strongly in cases with childhood stress. This is consistent with rodent studies, in which early-life and juvenile stress has been associated with persistent losses in dendritic spines in the PFC [27, 50], amygdala [50], and hippocampus [51], regions involved in regulating the stress response. These dendritic spine losses in the PFC were specific to young rodents, and not middle-aged or aged rodents [52]. In human imaging studies, exposure to early-life stress correlated with reduced grey matter volume [11] and processing speed [53] in the OFC, suggesting that underlying circuitry is altered by early-life stress. In support, children exposed to early-life stress are sensitised to threat processing, with increased perception of threats [54] and are more likely to have adverse behavioural responses [55]. Increased bias towards the perception of stress and impaired emotional/behavioural regulation associated with childhood stress have been hypothesised to underlie impaired developmental trajectories in neural circuits, in turn contributing to psychopathology [4, 56]. This suggests that exposure to stress in these vulnerable stages may dysregulate OFC circuits, contributing to altered emotional states [7, 8] and social behaviour [10] that might promote psychopathological development. Our data suggests these changes could be driven by persistent losses of dendritic spines on pyramidal neurons in the OFC.

We observed that mature mushroom spines were selectively affected by stress. Evidence for similar morphology-specific alterations in the rodent OFC is limited. This is further restricted by the lack of homology between the rodent OFC and human OFC which makes comparison difficult, regardless [19]. However, in the adjacent mPFC of rodents, reduced spine density in response to chronic psychological stress has been associated with a selective loss of thin spines, but this was recoverable after 21 days [52]. Similarly immediate, but recoverable, losses in thin spines were identified in the mPFC, although long-term reductions in mushroom spine head size and density have been noted [28, 57, 58]. In contrast, we identified that thin spine density was negatively correlated with distance from the soma, however, saw no significant losses in density. Morphology-specific alterations of mushroom spines due to psychosocial stress have also been identified in the hippocampus, and were associated with depression-like symptoms [51]. In a postmortem post-traumatic stress disorder cohort, selective losses of mushroom spines in the OFC were associated with increased FK506 binding protein 51 [32], a protein repeatedly associated with psychopathology in the presence of early-life stress [59]. There is a strong evidence that long-term remodelling of dendritic spines persistently affects mushroom spines in regions involved in stress regulation. It is important to note that although mushroom spines represent mature morphologies, this does not imply that other morphologies are inherently immature or not functional [60]. The current findings do, however, suggest that a closer exploration of how stress influences human spine dynamics may be important to resolving a role for stress in psychopathology.

Observed spine-specific changes were more evident in the superficial layers of the cortex. Cortical dendrites in the superficial layers appear to be particularly sensitive to stress, displaying selective reduced dendritic arborage [61] and spine density [37, 43, 44, 62]. These lamina-specific effects on synapses are also seen in the cortex in cases of both schizophrenia and depression [49, 63, 64]. Neurons in layers II/III of the OFC are involved in cortical signalling [65] including inputs from several sensory cortices (visual, somatosensory, olfactory and gustatory) [66]. These circuits are closely involved with valuing, processing, and responding to stimuli [7] in both cognitive [67] and affective [68] roles. Decline of these functions is associated with the development of psychopathology including depressive states and psychosis [69, 70]. In accordance, cortical circuit dysregulation has been identified in functional magnetic resonance imaging studies of psychiatric disorders specifically associated with childhood stress [71, 72]. Taken together, reduced spines in the OFC could be involved in impaired emotional regulation and anxiety associated with psychiatric disorders. This suggests that stress history is an important facilitator in loss of top-down control commonly seen in psychiatric disorders, and these circuits are particularly sensitive to the effects of stress during childhood.

We also evaluated the impact of stress on GR levels in the OFC, and the relationship with dendritic spine morphology. While we detected no difference in GR mRNA or protein between stress groups, GR protein negatively correlated with total and thin spine density, and GR mRNA levels negatively correlated to mushroom spine density. Decreases to GR mRNA/protein were previously identified in the OFC in bipolar disorder and schizophrenia [73]. In rodents, blocking GR using the antagonist mifepristone impaired cortical dendritic spine formation in non-stressed animals [41, 74]. GR is involved in upregulating spine number and maturation [40, 75], however, excess GR reduces spine density [76] and may contribute to towards the observed negative correlations. Given that GR has been located in the PSD [39], it is likely that alterations to GR levels are associated with specific effects on mushroom spines and, to a lesser extent, thin spines.

Study limitations include that postmortem analysis cannot capture individual fluctuations in stress exposure, symptom severity or progression which likely affect individual connectivity and dendritic spine populations. Given dendritic spines display age-associated reductions in density, older cohorts may under-represent the variance of spine alterations [77]. However, controlling for pH, RIN, PMI, and age is an effective measure to limit underlying effects that may shape spine populations [78]. Furthermore, although pH significantly varied between groups in our cohort (Table 1), inclusion of pH as a covariate did not alter the significance of our findings.

Another limitation lies in the Golgi staining process itself. Although this technique is widely utilised [79], it likely under-represents smaller spine morphologies [80]. This is particularly relevant as there is rodent studies suggest that smaller spine types are distinctly vulnerable to stress [52]. In accordance, the quantification technique only allowed for the identification of spines in the same x-y plane as the dendrite. Although the densities in the present study are comparable to those using similar methods [32], these combined factors likely underlie why the density of spines in the present study is lower than that identified using more sensitive methods in both rodents [81] and humans [82]. In the interpretation of the presented data, we acknowledge that these impacts may actually under-represent spine losses in the human OFC and the effect size may indeed be larger using techniques which facilitate greater resolution of spine detection.

Of note, we did not have access to subjects who had experienced severe stress, but did not develop psychopathology. Inclusion of this group when relevant patient samples become available will be helpful to separate the effects of stress from the effects of psychopathology. We hypothesise that non-psychiatric subjects exposed to severe stress may exhibit less pronounced or less persistent alterations of dendritic spines compared to those who exhibit psychopathological symptoms, which may be reflective of recovery and/or resilience mechanisms. This is an important avenue of exploration given the emerging role of the OFC in mediating stress resilience [83].

This study shows strong associations of stress exposure with persistent changes to spine density, particularly mushroom spines. Our findings suggest that OFC-related neuropathology in psychiatric disorders is differentially affected by severe stress, particularly if experienced early in life, and that these effects may be lifelong. Further studies to understand the molecular cascade associated with dendrite spine loss on pyramidal cells, and what this means for cortical cytoarchitecture and circuitry, is an important way forward.

## Acknowledgements

Dr Matosin was supported by an NHMRC Early Career Fellowship (APP1105445) and grants from the Brain Behaviour Research Foundation (NARSAD Young Investigator Grant, #26486) and the Rebecca L. Cooper Medical Research Foundation (#PG2020645). Dr Mechawar’s research is supported by grants from CIHR, HBHL and ERA-NET NEURON. Dr Binder’s research is supported by grants from NIMH, BMBF, EU-Horizon 2020 and the Hope for Depression Research Foundation. Tissues were received from the New South Wales Brain Tissue Resource Centre at the University of Sydney which is supported by the University of Sydney. Research reported in this publication was supported by the National Institute of Alcohol Abuse and Alcoholism of the National Institutes of Health under Award Number NIAAA012725-15. The content is solely the responsibility of the authors and does not represent the official views of the National Institutes of Health.

## References

[1] A. Schmitt, B. Malchow, A. Hasan, P. Falkai, The impact of environmental factors in severe psychiatric disorders Front Neurosci 8 (2014) 19.

[2] D. Vigo, G. Thornicroft, R. Atun, Estimating the true global burden of mental illness, Lancet Psychiat 3(2) (2016) 171–178.

[3] B.S. McEwen, P.J. Gianaros Stress-and allostasis-induced brain plasticity, Annu Rev Med 62 (2011) 431–445.

[4] W.E. Copeland, L. Shanahan, J. Hinesley, R.F. Chan, K.A. Aberg, J.A. Fairbank, et al., Association of childhood trauma exposure With adult psychiatric disorders and functional outcomes JAMA Netw Open 1(7) (2018) e184493.

[5] D.A. Glei, N. Goldman, Y.L. Chuang, M. Weinstein, Do chronic stressors lead to physiological dysregulation? Testing the theory of allostatic load Psychosomatic medicine 69(8) (2007) 769–776.

[6] K.E. Stephan, D.R. Bach, P.C. Fletcher, J. Flint, M.J. Frank, K.J. Friston, et al., Charting the landscape of priority problems in psychiatry, part 1: classification and diagnosis Lancet Psychiat 3(1) (2016) 77–83.

[7] M.L. Kringelbach, E.T. Rolls, The functional neuroanatomy of the human orbitofrontal cortex: evidence from neuroimaging and neuropsychology Prog Neurobiol 72(5) (2004) 341–372.

[8] A. Hahn, P. Stein, C. Windischberger, A. Weissenbacher, C. Spindelegger, E. Moser, et al., Reduced resting-state functional connectivity between amygdala and orbitofrontal cortex in social anxiety disorder NeuroImage 56(3) (2011) 881–889.

[9] N.H. Kalin, S.E. Shelton, R.J. Davidson, Role of the Primate Orbitofrontal Cortex in Mediating Anxious Temperament Biol Psychiatry 62(10) (2007) 1134–1139.

[10] J. Hornak, J. Bramham, E.T. Rolls, R.G. Morris, J. O’Doherty, P.R. Bullock, et al., Changes in emotion after circumscribed surgical lesions of the orbitofrontal and cingulate cortices Brain 126(Pt 7) (2003) 1691–1712.

[11] J.L. Hanson, M.K. Chung, B.B. Avants, E.A. Shirtcliff, J.C. Gee, R.J. Davidson, et al., Early stress is associated with alterations in the orbitofrontal cortex: a tensor-based morphometry investigation of brain structure and behavioral risk J Neurosci 30(22) (2010) 7466–7472.

[12] W.C. Drevets, Orbitofrontal cortex function and structure in depression, Annals of the New York Academy of Sciences 1121 (2007) 499–527.

[13] N. Kanahara, Y. Sekine, T. Haraguchi, Y. Uchida, K. Hashimoto, E. Shimizu, et al., Orbitofrontal cortex abnormality and deficit schizophrenia Schizophrenia research 143(2-3) (2013) 246–152.

[14] R. Mychasiuk, A. Muhammad, B. Kolb, Chronic stress induces persistent changes in global DNA methylation and gene expression in the medial prefrontal cortex, orbitofrontal cortex, and hippocampus Neuroscience 322 (2016) 489–499.

[15] C. Liston, M.M. Miller, D.S. Goldwater, J.J. Radley, A.B. Rocher, P.R. Hof, et al., Stress-induced alterations in prefrontal cortical dendritic morphology predict selective impairments in perceptual attentional set-shifting J Neurosci 26(30) (2006) 7870–7874.

[16] J. Bock, R.P. Murmu, N. Ferdman, M. Leshem, K. Braun, Refinement of dendritic and synaptic networks in the rodent anterior cingulate and orbitofrontal cortex: critical impact of early and late social experience Developmental neurobiology 68(5) (2008) 685–695.

[17] Z. Varga, D. Csabai, A. Miseta, O. Wiborg, B. Czéh, Chronic stress affects the number of GABAergic neurons in the orbitofrontal cortex of rats Behav Brain Res 316 (2017) 104–114.

[18] J.J. Miguel-Hidalgo, M. Moulana, P.H. Deloach, G. Rajkowska, Chronic Unpredictable Stress Reduces Immunostaining for Connexins 43 and 30 and Myelin Basic Protein in the Rat Prelimbic and Orbitofrontal Cortices Chronic Stress (Thousand Oaks) 2 (2018) 2470547018814186.

[19] J.D. Wallis, Cross-species studies of orbitofrontal cortex and value-based decision-making, Nature neuroscience 15(1) (2011) 13–19.

[20] H.B. Uylings, C.G. van Eden, Qualitative and quantitative comparison of the prefrontal cortex in rat and in primates, including humans Progress in brain research 85 (1990) 31–62.

[21] G. Eyal, M.B. Verhoog, G. Testa-Silva, Y. Deitcher, R. Benavides-Piccione, J. DeFelipe, et al., Human Cortical Pyramidal Neurons: From Spines to Spikes via Models Front Cell Neurosci 12(181) (2018).

[22] J. Herms, M.M. Dorostkar, Dendritic spine pathology in neurodegenerative diseases Annu Rev Pathol 11 (2016) 221–250.

[23] E.A. Nimchinsky, B.L. Sabatini, K. Svoboda, Structure and function of dendritic spines Annual review of physiology 64 (2002) 313–353.

[24] K.P. Berry, E. Nedivi, Spine Dynamics: Are They All the Same? Neuron 96(1) (2017) 43–55.

[25] K.-O. Lai, N.Y. Ip, Structural plasticity of dendritic spines: The underlying mechanisms and its dysregulation in brain disorders Biochim Biophys Acta 1832(12) (2013) 2257–2263.

[26] C. Yang, Y. Shirayama, J.-c. Zhang, Q. Ren, K. Hashimoto, Regional differences in brain-derived neurotrophic factor levels and dendritic spine density confer resilience to inescapable stress Int J Neuropsychopharmacol 18(7) (2015) 121.

[27] C. Helmeke, K. Seidel, G. Poeggel, T.W. Bredy, A. Abraham, K. Braun, Paternal deprivation during infancy results in dendrite-and time-specific changes of dendritic development and spine formation in the orbitofrontal cortex of the biparental rodent Octodon degus Neuroscience 163(3) (2009) 790–798.

[28] S.L. Gourley, A.M. Swanson, A.J. Koleske, Corticosteroid-induced neural remodeling predicts behavioral vulnerability and resilience J Neurosci 33(7) (2013) 3107–3112.

[29] H. Qiao, M.-X. Li, C. Xu, H.-B. Chen, S.-C. An, X.-M. Ma, Dendritic spines in depression: what we learned from animal models Neural Plast 2016 (2016) 8056370.

[30] C. Toro, J. Deakin, NMDA receptor subunit NRI and postsynaptic protein PSD-95 in hippocampus and orbitofrontal cortex in schizophrenia and mood disorder, Schizophrenia research 80(2-3) (2005) 323–330.

[31] A. Berdenis van Berlekom, C.H. Muflihah, G.J.L.J. Snijders, H.D. MacGillavry, J. Middeldorp, E.M. Hol, et al., Synapse pathology in schizophrenia: A meta-analysis of postsynaptic elements in postmortem brain studies Schizophr Bull 46(2) (2019) 374–386.

[32] K.A. Young, P.M. Thompson, D.A. Cruz, D.E. Williamson, L.D. Selemon, BA11 FKBP5 expression levels correlate with dendritic spine density in postmortem PTSD and controls Neurobiol Stress 2 (2015) 67–72.

[33] G. Turecki, M.J. Meaney, Effects of the social environment and stress on glucocorticoid receptor gene methylation: a systematic review Biol Psychiatry 79(2) (2016) 87–96.

[34] L.J. van der Knaap, H. Riese, J.J. Hudziak, M.M.P.J. Verbiest, F.C. Verhulst, A.J. Oldehinkel, et al., Glucocorticoid receptor gene (NR3C1) methylation following stressful events between birth and adolescence. The TRAILS study Transl Psychiatry 4(4) (2014) e381.

[35] F. Xiong, L. Zhang, Role of the hypothalamic-pituitary-adrenal axis in developmental programming of health and disease Front Neuroendocrinol 34(1) (2013) 27–46.

[36] J.M. McKlveen, B. Myers, J.N. Flak, J. Bundzikova, M.B. Solomon, K.B. Seroogy, et al., Role of prefrontal cortex glucocorticoid receptors in stress and emotion Biol Psychiatry 74(9) (2013) 672–679.

[37] M. Arango-Lievano, C. Peguet, M. Catteau, M.-L. Parmentier, S. Wu, M.V. Chao, et al., Deletion of neurotrophin signaling through the glucocorticoid receptor pathway causes tau neuropathology Sci Rep 6(1) (2016) 37231.

[38] M. Yoshiya, Y. Komatsuzaki, M. Ikeda, Y. Hojo, H. Mukai, Y. Hatanaka, et al., Corticosterone rapidly increases thorns of CA3 neurons via synaptic/extranuclear glucocorticoid receptor in rat hippocampus Front Neural Circuits 7 (2013) 191.

[39] L.R. Johnson, C. Farb, J. Morrison, B. McEwen, J. LeDoux, Localization of glucocorticoid receptors at postsynaptic membranes in the lateral amygdala Neuroscience 136(1) (2005) 289–299.

[40] Y. Komatsuzaki, Y. Hatanaka, G. Murakami, H. Mukai, Y. Hojo, M. Saito, et al., Corticosterone induces rapid spinogenesis via synaptic glucocorticoid receptors and kinase networks in hippocampus PloS one 7(4) (2012) e34124.

[41] C. Liston, W.-B. Gan, Glucocorticoids are critical regulators of dendritic spine development and plasticity in vivo, Proc Natl Acad Sci U S A 108(38) (2011) 16074–16079.

[42] C. von Economo, G.N. Koskinas, Atlas of Cytoarchitectonics of the Adult Human Cerebral Cortex 1st English ed. ed., Karger, Basil, New York, 2008.

[43] J.J. Radley, A.B. Rocher, M. Miller, W.G.M. Janssen, C. Liston, P.R. Hof, et al., Repeated stress induces dendritic spine loss in the rat medial prefrontal cortex Cereb Cortex 16(3) (2005) 313–320.

[44] R.M. Anderson, R.M. Glanz, S.B. Johnson, M.M. Miller, S.A. Romig-Martin, J.J. Radley, Prolonged corticosterone exposure induces dendritic spine remodeling and attrition in the rat medial prefrontal cortex J Comp Neurol 524(18) (2016) 3729–3746.

[45] J. Schindelin, I. Arganda-Carreras, E. Frise, V. Kaynig, M. Longair, T. Pietzsch, et al., Fiji: an open-source platform for biological-image analysis Nat Methods 9(7) (2012) 676–682.

[46] W.C. Risher, T. Ustunkaya, J. Singh Alvarado, C. Eroglu, Rapid Golgi analysis method for efficient and unbiased classification of dendritic spines PloS one 9(9) (2014) e107591.

[47] K.J. Livak, T.D. Schmittgen, Analysis of relative gene expression data using real-time quantitative PCR and the 2−ΔΔCT method Methods 25(4) (2001) 402–408.

[48] M.P. Forrest, E. Parnell, P. Penzes, Dendritic structural plasticity and neuropsychiatric disease Nat Rev Neurosci 19(4) (2018) 215–234.

[49] L.A. Glantz, D.A. Lewis, Decreased dendritic spine density on prefrontal cortical pyramidal neurons in schizophrenia Archives of general psychiatry 57(1) (2000) 65–73.

[50] C. Pinzón-Parra, B. Vidal-Jiménez, I. Camacho-Abrego, A.A. Flores-Gómez, A. Rodríguez-Moreno, G. Flores, Juvenile stress causes reduced locomotor behavior and dendritic spine density in the prefrontal cortex and basolateral amygdala in Sprague-Dawley rats, Synapse 73(1) (2019) e22066.

[51] S.D. Iñiguez, A. Aubry, L.M. Riggs, J.B. Alipio, R.M. Zanca, F.J. Flores-Ramirez, et al., Social defeat stress induces depression-like behavior and alters spine morphology in the hippocampus of adolescent male C57BL/6 mice Neurobiol Stress 5 (2016) 54–64.

[52] E.B. Bloss, W.G. Janssen, D.T. Ohm, F.J. Yuk, S. Wadsworth, K.M. Saardi, et al., Evidence for reduced experience-dependent dendritic spine plasticity in the aging prefrontal cortex J Neurosci 31(21) (2011) 7831–7839.

[53] A. Saleh, G.G. Potter, D.R. McQuoid, B. Boyd, R. Turner, J.R. MacFall, et al., Effects of early life stress on depression, cognitive performance and brain morphology Psychol Med 47(1) (2017) 171–181.

[54] K.A. McLaughlin, M.A. Sheridan, A.L. Gold, A. Duys, H.K. Lambert, M. Peverill, et al., Maltreatment exposure, brain structure, and fear conditioning in children and adolescents Neuropsychopharmacology 41(8) (2016) 1956–64.

[55] J.E. Shackman, S.D. Pollak, Impact of physical maltreatment on the regulation of negative affect and aggression Dev Psychopathol 26(4 Pt 1) (2014) 1021–33.

[56] K.A. McLaughlin, H.K. Lambert, Child trauma exposure and psychopathology: mechanisms of risk and resilience Curr Opin Psychol 14 (2017) 29–34.

[57] K.A. Michelsen, D.L. Van Den Hove, C. Schmitz, O. Segers, J. Prickaerts, H.W. Steinbusch, Prenatal stress and subsequent exposure to chronic mild stress influence dendritic spine density and morphology in the rat medial prefrontal cortex BMC Neuroscience 8(1) (2007) 107.

[58] J.J. Radley, A.B. Rocher, A. Rodriguez, D.B. Ehlenberger, M. Dammann, B.S. McEwen, et al., Repeated stress alters dendritic spine morphology in the rat medial prefrontal cortex J Comp Neurol 507(1) (2008) 1141–1150.

[59] N. Matosin, T. Halldorsdottir, E.B. Binder, Understanding the molecular mechanisms underpinning gene by environment interactions in psychiatric disorders: the FKBP5 model Biol Psychiatry 83(10) (2018) 821–830.

[60] M. Adrian, R. Kusters, C.J. Wierenga, C. Storm, C.C. Hoogenraad, L.C. Kapitein, Barriers in the brain: resolving dendritic spine morphology and compartmentalization Frontiers in Neuroanatomy 8 (2014).

[61] R.M. Shansky, C. Hamo, P.R. Hof, B.S. McEwen, J.H. Morrison, Stress-induced dendritic remodeling in the prefrontal cortex is circuit specific Cereb Cortex 19(10) (2009) 2479–2484.

[62] J.J. Radley, C.M. Arias, P.E. Sawchenko, Regional differentiation of the medial prefrontal cortex in regulating adaptive responses to acute emotional stress J Neurosci 26(50) (2006) 12967–12976.

[63] N. Kolluri, Z. Sun, A.R. Sampson, D.A. Lewis, Lamina-specific reductions in dendritic spine density in the prefrontal cortex of subjects with schizophrenia The American journal of psychiatry 162(6) (2005) 1200–1202.

[64] H.J. Kang, B. Voleti, T. Hajszan, G. Rajkowska, C.A. Stockmeier, P. Licznerski, et al., Decreased expression of synapse-related genes and loss of synapses in major depressive disorder Nat Med 18(9) (2012) 1413.

[65] C.R. Gerfen, M.N. Economo, J. Chandrashekar, Long distance projections of cortical pyramidal neurons J Neurosci Res 96(9) (2018) 1467–1475.

[66] D. Ongür, J.L. Price, The organization of networks within the orbital and medial prefrontal cortex of rats, monkeys and humans Cereb Cortex 10(3) (2000) 206–19.

[67] S. McClure, D. Laibson, G. Loewenstein, J. Cohen, Separate Neural Systems Value Immediate and Delayed Monetary Rewards Science 306(565) (2004) 503–507.

[68] A. Bechara, H. Damasio, A.R. Damasio, G.P. Lee, Different contributions of the human amygdala and ventromedial prefrontal cortex to decision-making J Neurosci 19(13) (1999) 5473–5481.

[69] W. Cheng, E.T. Rolls, J. Qiu, W. Liu, Y. Tang, C.-C. Huang, et al., Medial reward and lateral non-reward orbitofrontal cortex circuits change in opposite directions in depression Brain 139(12) (2016) 3296–3309.

[70] C. Fahim, E. Stip, A. Mancini-Marïe, S. Potvin, D. Malaspina, Orbitofrontal dysfunction in a monozygotic twin discordant for postpartum affective psychosis: a functional magnetic resonance imaging study Bipolar Disord 9(5) (2007) 541–5.

[71] M. Yu, K.A. Linn, R.T. Shinohara, D.J. Oathes, P.A. Cook, R. Duprat, et al., Childhood trauma history is linked to abnormal brain connectivity in major depression Proc Natl Acad Sci U S A 116(17) (2019) 8582–8590.

[72] S. Kim, J.S. Kim, M. Shim, C.-H. Im, S.-H. Lee, Altered cortical functional network during behavioral inhibition in individuals with childhood trauma Sci Rep 8(1) (2018) 10123–10123.

[73] D. Sinclair, M.J. Webster, J.M. Fullerton, C.S. Weickert, Glucocorticoid receptor mRNA and protein isoform alterations in the orbitofrontal cortex in schizophrenia and bipolar disorder BMC Psychiatry 12(1) (2012) 84.

[74] M. Pedrazzoli, M. Losurdo, G. Paolone, M. Medelin, L. Jaupaj, B. Cisterna, et al., Glucocorticoid receptors modulate dendritic spine plasticity and microglia activity in an animal model of Alzheimer’s disease Neurobiol Dis 132 (2019) 104568.

[75] M.F. Russo, S.R. Ah Loy, A.R. Battle, L.R. Johnson, Membrane associated synaptic mineralocorticoid and glucocorticoid receptors are rapid regulators of dendritic spines Front Cell Neurosci 10 (2016) 161–161.

[76] C. Liston, J.M. Cichon, F. Jeanneteau, Z. Jia, M.V. Chao, W.B. Gan, Circadian glucocorticoid oscillations promote learning-dependent synapse formation and maintenance Nature neuroscience 16(6) (2013) 698–705.

[77] M.L. Feldman, C. Dowd, Loss of dendritic spines in aging cerebral cortex Anat Embryol 148(3) (1975) 279–301.

[78] A.D. Stan, S. Ghose, X.-M. Gao, R.C. Roberts, K. Lewis-Amezcua, K.J. Hatanpaa, et al., Human postmortem tissue: what quality markers matter? Brain Res 1123(1) (2006) 1–11.

[79] G. Das, K. Reuhl, R. Zhou, The Golgi-Cox method Methods in molecular biology (Clifton, N.J.) 1018 (2013) 313–21.

[80] J.J. Mancuso, Y. Chen, X. Li, Z. Xue, S.T.C. Wong, Methods of dendritic spine detection: from Golgi to high-resolution optical imaging Neuroscience 251 (2013) 129–140.

[81] A.J. Whyte, H.W. Kietzman, A.M. Swanson, L.M. Butkovich, B.R. Barbee, G.J. Bassell, et al., Reward-Related Expectations Trigger Dendritic Spine Plasticity in the Mouse Ventrolateral Orbitofrontal Cortex The Journal of Neuroscience 39(23) (2019) 4595–4605.

[82] S.C. Das, D. Chen, W.B. Callor, E. Christensen, H. Coon, M.E. Williams, DiI-mediated analysis of presynaptic and postsynaptic structures in human postmortem brain tissue J Comp Neurol 527(18) (2019) 3087–3098.

[83] F. Kong, X. Ma, X. You, Y. Xiang, The resilient brain: psychological resilience mediates the effect of amplitude of low-frequency fluctuations in orbitofrontal cortex on subjective well-being in young healthy adults Soc Cogn Affect Neurosci 13(7) (2018) 755–763.

