## Supplemental Materials for "Severe childhood and adulthood stress associates with neocortical layer-specific reductions of mature spines in psychiatric disorders"

### Supplementary Materials

**Supplementary Table 1.** Detailed cohort demographics. Patient severe stress history was derived from medical records, following extensive contact of patients with healthcare professionals.

| Age | Sex | Classification | PMI | Brain pH | RIN | COD category | Stress exposures extracted from clinical notes |
|---|---|---|---|---|---|---|---|
| **Cases with childhood stress** | | | | | | | |
| **59** | Male | Schizophrenia-Paranoid | 48.5 | 6.56 | 7.8 | Cardiac | Childhood marked with domestic violence & 'put downs' from his father.<br>Mother died when he was 16. Father died when he was 19.<br>No family support except his sister, with whom he chose to have no contact. |
| **30** | Male | Schizophrenia-Paranoid | 37 | 6.32 | 5.9 | Toxicity | Early childhood demonstrated verbal and physical abuse by father<br>Parents divorced when donor was 18.<br>Donor did not cope with divorce well. |
| **54** | Male | Schizophrenia-Paranoid | 64 | 6.77 | 7.8 | Cardiac/Hepatic | Childhood had abusive and violent father<br>No friends. Always a loner and isolated. |
| **84** | Male | Schizophrenia-Disorganised | 5.5 | 6.55 | 5.2 | | Childhood upbringing 'disturbed' violence and sexual abuse by father.<br>Wife and child left him mid 20's. No further contact ever made with his child. |
| **83** | Female | Schizophrenia | 23 | 6.23 | 5.5 | Renal | Early childhood physical and sexual abuse by father.<br>Witnessed physical and sexual abuse to other family members.<br>Parents separated at age 16 |
| **74** | Male | Major Depression | 39.5 | 6.59 | 6.6 | Respiratory/Toxicity | At age 7 was sent away to boarding school.<br>Abuse and neglect from parents.<br>Severe depressive episodes until death. |
| **58** | Female | Major Depression | 39 | 6.02 | 4.4 | Trauma | Sexually assaulted at age 5.<br>4 suicide attempts. |
| **34** | Male | Schizoaffective | 26 | 6.7 | 8.3 | Hanging | Poor childhood.<br>Poor nutrition.<br>Severe depression.<br>Suicide attempts |
| **Cases with adult stress** | | | | | | | |
| **34** | Male | Bipolar Disorder | 68 | 6.72 | 8.7 | Toxicity | Did not cope well with mother's death when donor was 31. |
| **48** | Male | Bipolar Disorder 1-Manic episode co-morbid alcohol abuse | 23.5 | 6.34 | 6.1 | Cardiac | Age 23 relationship breakdown. Age 38 marriage breakdown. Access to children difficult |
| **60** | Female | Major Depression | 28.5 | 6.61 | 6.2 | Trauma | Discharged from Army aged 46. Severe depression with suicidal ideation. Cause of death suicide. |

| | | | | | | | |
|---|---|---|---|---|---|---|---|
| **33** | Male | Schizoaffective Disorder- Bipolar type | 48 | 6.65 | 8.2 | Hanging | Aged 20 mental health deteriorated. Parents split when donor in his twenties. Many jobs, unable to maintain any. Convictions for stealing & driving. |
| **61** | Female | Schizophrenia | 60.5 | 5.84 | 2.9 | Metabolic | Age 27 lost her first-born child. Second child left the family home & chose to live with grandmother at age 15 instead of mother – this caused much distress. |
| **56** | Male | Schizophrenia-Paranoid | 65 | 6.56 | 8 | Respiratory | At age 30 donor had major motor vehicle accident and begun to drink heavily. Married briefly for a few months. Daughter from that relationship. Never kept contact with daughter. |
| **64** | Male | Schizophrenia-Paranoid | 48 | 6.66 | 8.1 | Cardiac | Extremely introverted. Mother died when donor was aged 23. Donor shot himself when he found out, requiring emergency surgery. |
| **63** | Male | Schizophrenia-Paranoid | 25 | 6.63 | 6.7 | Cardiac | Mother died and behaviour and mood subsequently deteriorated significantly. |
| **Cases no severe stress** | | | | | | | |
| **60** | Female | Schizophrenia | 27.5 | 6.44 | 7.3 | Cardiovascular | No trauma |
| **49** | Male | Schizophrenia chronic-continuous | 22 | 6.33 | 6.1 | Respiratory | No trauma |
| **40** | Female | Schizophrenia-Undifferentiated | 20 | 6.63 | 7.6 | Respiratory | No trauma |
| **50** | Male | Schizophrenia-Undifferentiated / Tumour | 42 | 6.71 | 8.6 | Cancer | No trauma |
| **52** | Male | Schizophrenia | 46 | 6.43 | 7.9 | Cardiac | No trauma |
| **31** | Female | Major Depression | 38 | 6.4 | 7 | Respiratory/Toxicity | No trauma |
| **64** | Male | Major Depression | 24 | 6.6 | 7.6 | Hanging | No trauma |
| **60** | Male | Bipolar Disorder | 33.5 | 6.6 | 6.7 | Respiratory/Toxicity | No trauma |
| **50.75** | 5M/3F | 5 SZ, 2 MDD, 1 BPD | 31.625 | 6.5175 | 7.35 | | |
| **Controls no severe stress** | | | | | | | |
| **33** | Female | Control | 24 | 6.77 | 7.7 | Cardiac | No trauma |
| **49** | Female | Control | 15 | 6.93 | 8.6 | Cardiac | No trauma |
| **69** | Female | Control | 39 | 6.72 | 7.9 | Cardiac/Respiratory | No trauma |
| **73** | Male | Control | 48 | 6.8 | 7.9 | Cardiac | No trauma |
| **62** | Male | Control | 37.5 | 6.56 | 9 | Cardiac | No trauma |
| **59** | Male | Control | 28 | 6.77 | 7.3 | Cardiac | No trauma |

| | | | | | | | |
|---|---|---|---|---|---|---|---|
| **32** | Male | Control | 36 | 6.77 | 7.9 | Cardiac | No trauma |
| **64** | Male | Control | 30 | 6.82 | 7.5 | Cardiac | No trauma |

**Supplementary Table 2.** Summary statistics glucocorticoid receptor mRNA/protein levels and total, mushroom, thin, stubby, and filopodia spine density in the total cohort (*n*=32) and within individual groups (*n*=8). Significance indicated by bold. Significant correlations denoted as adjusted R-squared.

| **Spine** | **Layer** | **Total cohort** | | **Control** | | **No Stress[1]** | | **Adulthood Stress[1]** | | **Childhood Stress[1]** | |
|---|---|---|---|---|---|---|---|---|---|---|---|
| | | mRNA | Protein | mRNA | Protein | mRNA | Protein | mRNA | Protein | mRNA | Protein |
| **Total** | Combined | T=2.015<br>P=0.0533 | **T=2.526<br>P=0.0170<br>$R^2$=0.1479** | T=1.415,<br>P=0.2069 | T=1.289<br>P=0.245 | T=1.365<br>P=0.2210 | T=1.330<br>P=0.232 | T=0.450<br>P=0.671 | T=1.321<br>P=0.235 | T=1.675<br>P=0.1449 | T=1.737<br>P=0.1330 |
| | II/III | T=0.0769<br>P=0.448 | T=1.418<br>P=0.1665 | T=1.501<br>P=0.1840 | T=1.399<br>P=0.211 | T=0.242<br>P=0.8170 | T=0.146<br>P=0.889 | T=1.075<br>P=0.3314 | T=1.082<br>P=0.321 | T=0.875<br>P=0.4155 | T=0.918<br>P=0.394 |
| | V | **T=2.328<br>P=0.0273<br>$R^2$=0.1323** | **T=2.724<br>P=0.01081<br>$R^2$=0.1763** | T=1.468<br>P=0.2020 | T=1.521<br>P=0.189 | T=1.664<br>P=0.1471 | T=1.727<br>P=0.1350 | T=0.023<br>P=0.9825 | T=1.411<br>P=0.208 | T=1.822<br>P=0.1183 | T=1.870<br>P=0.1107 |
| **Mushroom** | Combined | **T=2.093<br>P=0.0452<br>$R^2$=0.1013** | T=1.047<br>P=0.3032 | T=1.825<br>P=0.118 | T=1.687<br>P=0.1425 | T=1.397<br>P=0.2119 | T=1.417<br>P=0.206 | T=0.708<br>P=0.5106 | T=0.152<br>P=0.884 | T=1.147<br>P=0.295 | T=1.179<br>P=0.2831 |
| | II/III | T=0.548<br>P=0.5880 | T=0.029<br>P=0.977 | T=0.269<br>P=0.7986 | T=0.305<br>P=0.772 | T=1.356<br>P=0.233 | T=1.379<br>P=0.226 | T=0.324<br>P=0.7624 | T=0.018<br>P=0.986 | T=0.020<br>P=0.9850 | T=0.010<br>P=0.993 |
| | V | T=1.250<br>P=0.2220 | T=0.473<br>P=0.6402 | T=0.686<br>P=0.5234 | T=0.672<br>P=0.531 | T=0.647<br>P=0.5417 | T=0.706<br>P=0.507 | T=0.141<br>P=0.8948 | T=0.814<br>P=0.452 | T=2.344<br>P=0.0661 | T=2.327<br>P=0.0674 |
| **Thin** | Combined | T=1.357<br>P=0.185 | **T=2.254<br>P=0.03167<br>$R^2$=0.1163** | T=0.703<br>P=0.5082 | T=0.609<br>P=0.565 | T=0.687<br>P=0.5178 | T=0.723<br>P=0.497 | T=0.925<br>P=0.3976 | **T=2.603<br>P=0.0405<br>$R^2$=0.4521** | T=0.574<br>P=0.5866 | T=0.601<br>P=0.564 |
| | II/III | T=0.572<br>P=0.572 | T=1.629<br>P=0.1138 | T=1.157<br>P=0.2911 | T=1.081<br>P=0.321 | T=0.142<br>P=0.8919 | T=0.168<br>P=0.872 | T=0.144<br>P=0.8910 | T=2.054<br>P=0.0857 | T=1.275<br>P=0.2495 | T=1.237<br>P=0.262 |
| | V | T=1.669<br>P=0.106 | **T=2.245<br>P=0.03229<br>$R^2$=0.1153** | T=0.042<br>P=0.9679 | T=0.052<br>P=0.960 | T=1.004<br>P=0.3541 | T=1.047<br>P=0.335 | T=1.209<br>P=0.2809 | T=2.077<br>P=0.0831 | T=1.754<br>P=0.1300 | T=1.802<br>P=0.1215 |
| **Stubby** | Combined | T= 0.787<br>P=0.438 | T=1.415<br>P=0.1675 | T=0.358<br>P=0.7323 | T=0.361<br>P=0.731 | T=0.374<br>P=0.7213 | T=0.294<br>P=0.779 | T=0.394<br>P=0.7099 | T=0.776<br>P=0.467 | **T=2.512<br>P=0.0458<br>$R^2$=0.4314** | **T=3.482<br>P=0.0131<br>$R^2$=0,4408** |
| | II/III | T=0.067<br>P=0.947 | T=0.635<br>P=0.530 | T=0.594<br>P=0.5744 | T=0.603<br>P=0.569 | T=0.087<br>P=0.994 | T=0.188<br>P=0.857 | T=1.021<br>P=0.3539 | T=0.339<br>P=0.746 | **T=2.465<br>P=0.0488<br>$R^2$=0.4202** | **T=2.535<br>P=0.0444<br>$R^2$=0.4367** |

| | | | | | | | | | | | |
|---|---|---|---|---|---|---|---|---|---|---|---|
| | V | T=0.1251<br>P=0.221 | T=1.749<br>P=0.0906 | T=0.138<br>P=0.8951 | T=0.144<br>P=0.890 | T=0.626<br>P=0.555 | T=0.599<br>P=0.571 | T=0.292<br>P=0.7812 | T=1.273<br>P=0.250 | T=1.311<br>P=0.238 | T=1.313<br>P=0.2371 |
| **Filopodia** | Combined | T=0.841<br>P=0.407 | T=1.147<br>P=0.260 | T=0.596<br>P=0.573 | T=0.634<br>P=0.550 | T=0.547<br>P=0.6044 | T=0.502<br>P=0.633 | T=0.672<br>P=0.5312 | T=0.855<br>P=0.425 | T=1.273<br>P=0.25 | T=1.242<br>P=0.261 |
| | II/III | T=1.453<br>P=0.157 | T=1.616<br>P=0.117 | T=1..43<br>P=0.203 | T=1.413<br>P=0.207 | T=1.156<br>P=0.2916 | T=1.162<br>P=0.289 | T=0.260<br>P=0.8055 | T=0.652<br>P=0.538 | T=0.193<br>P=0.853 | T=0.192<br>P=0.854 |
| | V | T=0.108<br>P=0.195 | T=0.174<br>P=0.863 | T=0.944<br>P=0.382 | T=0.850<br>P=0.428 | T=0.148<br>P=0.8869 | T=0.212<br>P=0.839 | T=0.736<br>P=0.4950 | T=0.485<br>P=0.645 | T=1.754<br>P=0.13 | T=1.701<br>P=0.140 |

[1] Consisting of cases of cross-diagnosis psychiatric disorders (major depressive disorder, bipolar disorder, schizophrenia, schizoaffective disorder).

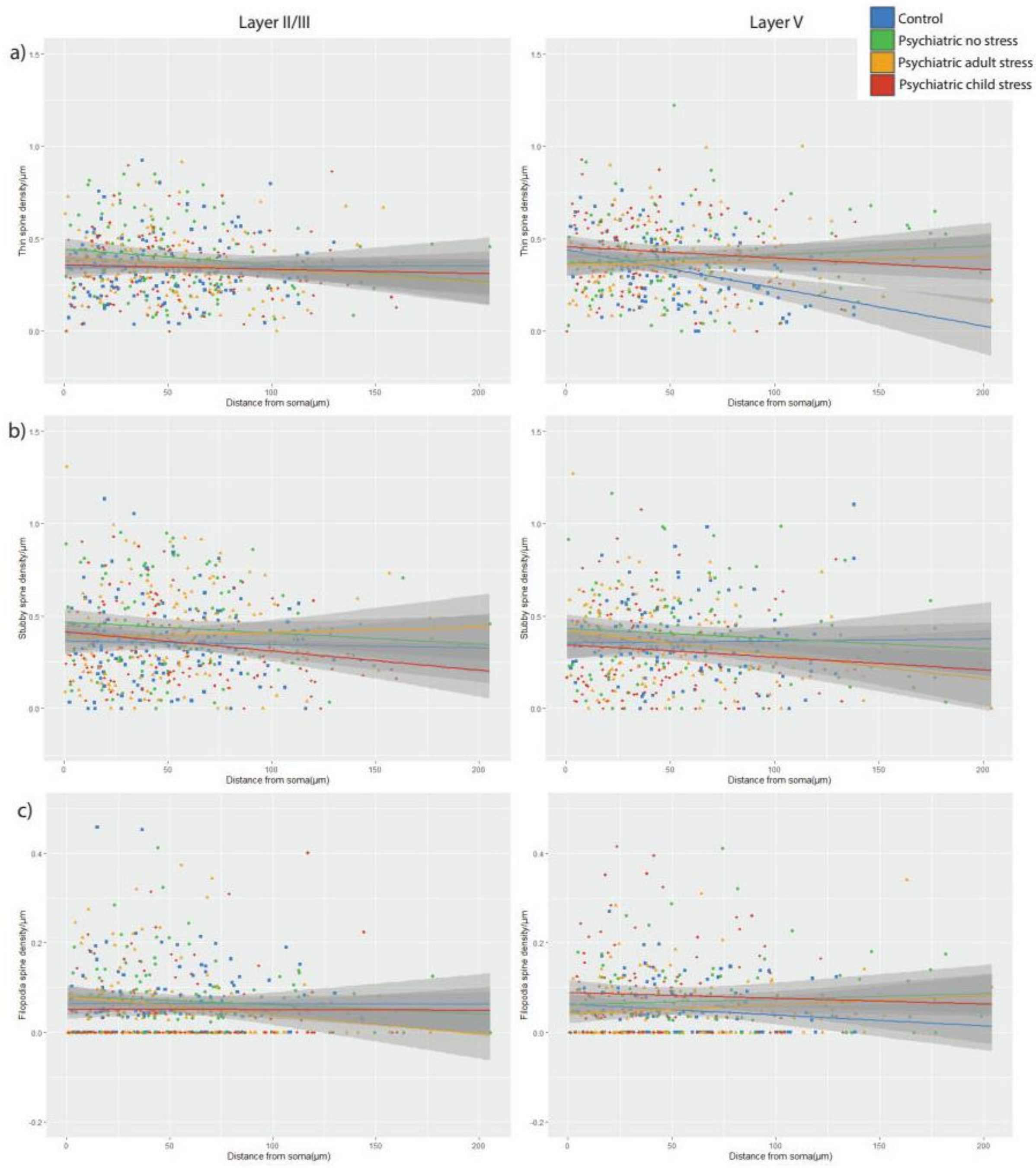

**Supplementary Figure 1.** Linear correlations (± confidence interval) between segment distance from the soma and (**A**) thin spine density (t=2.063, P=0.0396, R2=0.0065) (**B**) stubby spine density, **C)** filopodia spine density, measured along the dendritic segment. No interactive effect between groups was identified.

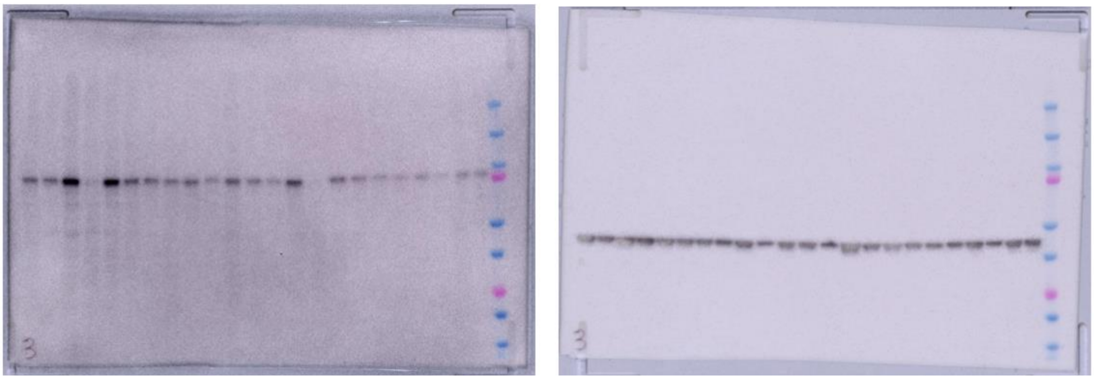

**Supplementary Figure 2.** Representative western blot showing a single band detected for both NR3C1 (left blot) and β-actin (right blot) at the expected molecular weights of 94 kDa and 42 kDa, respectively.
